# Targeting endogenous retrovirus gene transcription in human cancers

**DOI:** 10.1101/449686

**Authors:** Audrey T. Lin, Cindy G. Santander, Emanuele Marchi, Timokratis Karamitros, Aris Katzourakis, Gkikas Magiorkinis

## Abstract

Endogenous retroviruses (ERVs) are remnants of ancient retroviral infections that make up to 8% of the human genome. Although these elements are mostly fragmented and inactive, many proviruses belonging to the HERV-K (HML-2) family, the only lineage still proliferating in the genome after the human-chimpanzee split, have intact open reading frames, some encoding for accessory genes called *np9* and *rec* that interact with oncogenic pathways. Many studies have established that the transient expression of ERVs are in both stem cells and cancers results in aberrant self-renewal and uncontrolled proliferation.

The wealth of high-quality genomic and transcriptomic Illumina sequence data available from The Cancer Genome Atlas (TCGA) that are sequenced from a diversity of different tumour types makes it a valuable resource in cancer research. However, there is currently no universal computational method for inferring expression of specific repetitive elements from RNA-seq data, such as genes encoded by HERV-K (HML-2).

This study presents a novel and a highly specific pipeline that is able to capture and measure transcription of *np9* and *rec* encoded by proviruses that share great sequence similarity, and are transcribed at very low levels. We show by using our novel methodology that *np9* and *rec* are overexpressed in breast cancer, germ cell tumours, skin melanoma, lymphoma, ovarian cancer, and prostate cancer compared to non-diseased tissues. We also show that *np9* and *rec* are specifically expressed in the 8 and 16-cell stage in human preimplantation embryos.

## Introduction

Endogenous retroviruses (ERVs) are remnants of ancient retroviral infections that make up about 5-8% of the human genome. These ancient elements are mostly inactive due to mutations or deletions, or are epigenetically repressed. Many proviruses belonging to HERV-K (HML-2) (shortened to HK2 here) family, the only lineage still proliferating in the genome after the human-chimpanzee split, have intact open reading frames, some encoding for accessory proteins called Np9 and Rec that have been found to interact with cellular pathways linked to cancer [1],[2],[3]. Type I proviruses have a 292 base pair deletion in *env* that leads to a different splice donor, producing an *np9* transcript instead of *rec*, which is encoded by type II proviruses [4],[5] (**Fig. S1**). Rec is a functional homolog of accessory proteins Rev and Rex, encoded by HIV-1 and T-cell leukaemia virus type 1 (HTLV-1) respectively. Rev/Rex/Rec functions in regulating viral gene expression by transporting viral mRNA from the nucleus into the cytosol [6],[7]. Rec protein was found to interfere with germ cell development and also cause carcinoma in mice [8].

It is unclear what function *np9* has in virus replication, if any, but it does appear to have oncogenic activity. Np9 protein can activate the ERK, Akt and Notch1 pathways, and upregulate β-catenin, promoting survival and growth of leukaemia stem/progenitor cells [1]. Np9 protein is also preferentially expressed in transformed cells and is known to interact with LNX, an important part of the Notch signal transduction pathway, implicated in regulating breast and prostate cancer proliferation [5],[3]. Its expression significantly promotes the growth of leukaemia cells, and when silenced, the growth of myeloid and lymphoblastic leukaemic cells is inhibited. Both Np9 and Rec proteins interact with promyelocytic leukaemia zinc finger (PLZF) tumour suppressor, a chromatin remodeller and transcriptional repressor that is implicated in spermatogonial stem cell renewal and cancer. The Np9-Rec-PLZF interaction results in the de-repression of the c-*myc* proto-oncogene, c-Myc overexpression, altered expression of genes regulated by c-Myc, and increased cell survival and proliferation [2]. Np9 protein is also reported to bind and regulate ubiquitin ligase MDM2, which represses the activity of p53 tumour suppressor [9]. Transcripts of *np9* and *rec* are also found in human embryonic stem cells and human induced pluripotent stem cell lines, and are associated with maintenance of pluripotency [10],[11].

With recent advances in high-throughput mRNA sequencing (RNA-seq), it is possible in a single assay to find novel genes and transcripts as well as measure transcription [12],[13]. Small RNA-seq experiments are capable of producing vast volumes of data: current sequencers can generate more than 500 gigabases of raw sequencing reads per run [14]. The sensitivity of RNA-seq allows for the potential detection of alternative splice isoforms of transcripts as well as rare and cell and context-specific transcripts. Moreover, because the number of reads that are produced from an RNA transcript is a function of the transcript abundance, the depth of read coverage, or read density, can be used as a proxy to measure gene expression [15],[16],[14]. However, there may be inherent biases in the RNA-seq methodology that must be taken into consideration. In general, most sequencing platforms produce short sequence reads that are assembled computationally to generate full-length transcripts. Even for fairly well-characterised transcripts, such as from protein-coding genes, ambiguity arises from the expression of transcriptional variants that often share exons.

Given the sensitivity of RNA-seq, it is likely that HERV transcripts can be detected, even if they are expressed in low abundance. However, because HERV transcripts are often longer than the lengths of reads generated by most sequencing platforms, and that HERVs are often greater than 97% identical to one another [17], there are many uncertainties arising from analysing HERVs in RNA-seq data. Due to the high similarity of sequences from repetitive regions, it is difficult to reconstruct these transcripts with accuracy. Problems with library preparation as well as GC content of the transcript also produce biases [18]. Inconsistent coverage can result from biological stochasticity – transcripts can be expressed at different levels and at different lengths depending on cellular conditions [19]. As a result, measuring the expression of the alternatively spliced *np9* and *rec* transcripts with confidence can be challenging.

To the best of our knowledge, there is currently no standardised computational method for inferring expression of repetitive elements from RNA-seq data. We designed a pipeline that enabled us to measure the cDNA transcripts of *np9* and *rec* in mapping RNA-seq reads to a reference transcript and only counting the reads spanning the splice junction. This ensures the measurement of these HERV alternative splice products with high sequence similarity with great specificity. By using our novel methodology of processing and analysis of publicly available and controlled-access RNA-seq data, we will show overexpression of the HK2 targets in the cancer types that are known in the literature to overexpress HERVs. The cancers we had expected to see HERV overexpression in comparison to non-diseased tissue include breast cancer, germ cell tumours, skin melanoma, lymphoma, ovarian cancer, and prostate cancer.

In addition, it had been reported that during early human embryogenesis, HK2 is transcribed only during the 8-cell, 16-cell (morula), in early blastocyst outgrowth stages, as seen in early passages of human embryonic stem cell (hESC) lines [7],[20]. In order to determine whether *np9* and *rec* follow these patterns of expression, published single cell RNA-seq data [20] was used to measure the transcription of *rec* and *np9* in human preimplantation embryos.

Using RNA-seq data from The Cancer Genome Atlas and other repositories, we observed high transcription of *np9* and *rec* in testicular germ cell cancer, and cancers of the eye, skin, liver, lung, and thymus, as well as during the 8- and 16-stage cell stages of human embryogenesis.

## Materials and Methods

### cDNA reference sequences

The 225 bp *np9* cDNA sequence (Genbank accession number JA719353.1) was used as a reference for the RNA-seq reads to map against. Because there are no other known isoforms of *np9*, no consensus sequence was created. However, if the splice junction is shorter than the 48bp-long RNA-seq read, the read will not map properly even if it spans the junction, because the read would be longer than the *np9* sequence leading up to the splice junction (48 bp vs 44 bp long, respectively) (**Fig. 2**).

To capture these reads that would otherwise be lost, 52 bp from the 3’ end of HERV-K (HML-2.HOM) *pol* (accession number AF074086.2) was added to the *np9* cDNA reference, forming a 274 bp sequence, allowing the full RNA-seq read to map to the *np9* reference (**Table S1**).

All *rec*, or complete ORF (cORF) as reported on Genbank sequences were downloaded from Genbank. Because multiple isoforms of *rec* transcript are known, including a 414 bp isoform with a 58 bp deletion containing the splice donor [21], sequences of the longer 471 bp isoform were downloaded and aligned, and a consensus sequence was created. All RNA-seq reads were mapped to this *rec* consensus sequence (**Table S1**).

*pol* transcript from HK2 proviruses and HERV-K113 5’ LTR (accession number NC_022518.1) transcript were used to quantify the background transcription level of HK2 loci in both solid primary tumour and solid non-diseased tissue samples. Full type 1 and type 2 HK2 provirus sequences were downloaded [17], aligned, and a consensus sequence created. A 532 bp fragment from the 5’ end of the *pol* consensus sequence was used to map all RNA-seq reads to (**Table S1**). All consensus sequences were created in Geneious version 8.1 [22].

### Alignment

Because of the polymorphic nature of HK2 integrations, it is unknown how many proviruses express *np9* and *rec* in the genome, from one individual to the next. Therefore, transcription levels within each sample were standardised by selecting a “housekeeping” or “reference” gene to normalise HK2 activity with *actb* (accession number NM_001101.3) was used as a reference gene in all the samples analysed.

The RNA-seq reads were aligned with bowtie2 v.2.2.5 [23] to all cDNA reference sequences in end-to-end mode, using default settings. All aligned reads in the resulting BAM file with a mapping quality of less than 20 were filtered out using Samtools version 1.2, [24] which was then followed by sorting and indexing with Picard Tools version 2.1.1 (Broad Institute), for visualization on Integrative Genome Browser (IGV) (Broad Institute).

### Filtering for np9 and rec-specific aligned reads

The high sequence identity between different HK2 proviruses necessitates stringent filtering to ensure that the reads mapped to the *np9* and *rec* reference sequences indeed belonged to expressed *np9* and *rec* transcripts. For this reason, the junction between the splice donor and the splice acceptor site was the critical target in capturing *np9* and *rec* transcripts within each given sample. In order to ensure high specificity when capturing alternative splice products of the same gene, within the filtered BAM file, all reads that did not span the splice junction of *np9* and *rec* within 40 bp in either direction were discarded. Also discarded were reads mapped to *np9* reference sequence with >1 mismatch, and >2 mismatches for *rec* (**Fig. 1, 2**). The mismatch thresholds differ to allow for capture of transcripts expressed from isoforms of *rec*.

**Figure 1.**
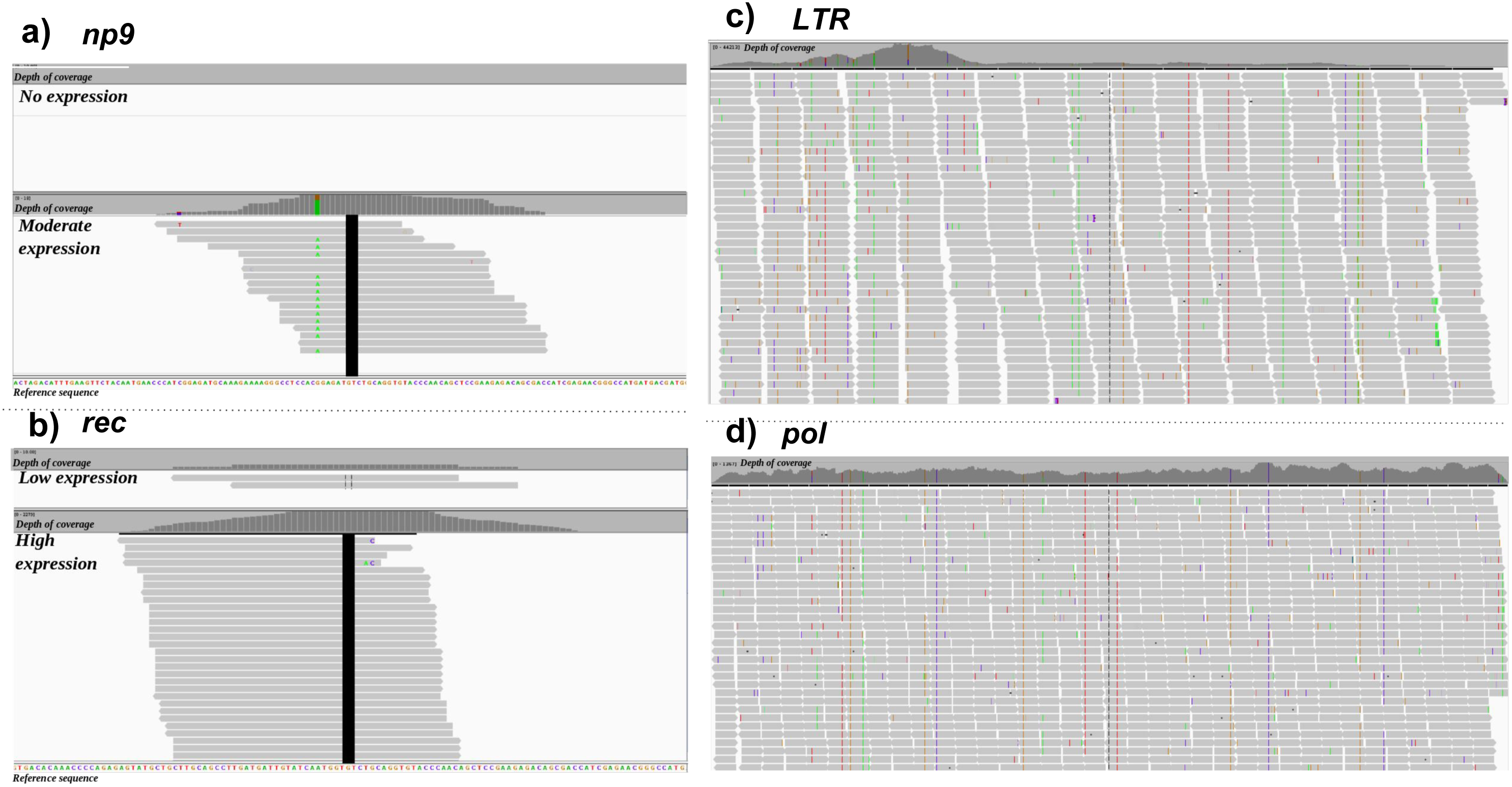
Sequenced cDNA reads from primary tumours mapped to HK2 targets as viewed on Integrative Genome Browser (IGV). The horizontal shaded boxes display the depth of coverage of mapped reads. Thick vertical black bars indicate the splice junction of *np9* and *rec.* A segment of the reference sequence is displayed at the bottom of each panel. (**a.)** RNA-seq reads from lung adenocarcinoma (LUAD) primary tumour samples from two different cancer patients mapped to *np9* reference sequence. The top panel shows a patient with no *np9* expression. The bottom panel shows a patient with moderate *np9* expression. **(b.)** RNA-seq reads from testicular germ cell tumour (TGCT) primary tumour samples from two different cancer patients mapped to *rec* reference sequence. The top panel shows a patient with low *rec* expression, with only 2 reads that map perfectly to the reference. The bottom panel shows a patient with high expression of *rec*. **(c.)** RNA-seq reads from testicular germ cell tumour (TGCT) primary tumour sample mapped to HERV-K113 LTR reference sequence. LTR is highly expressed. **(d.)** RNA-seq reads from testicular germ cell tumour (TGCT) primary tumour sample mapped to HK2 *pol* reference sequence. *pol* is highly expressed.

**Figure 2.**
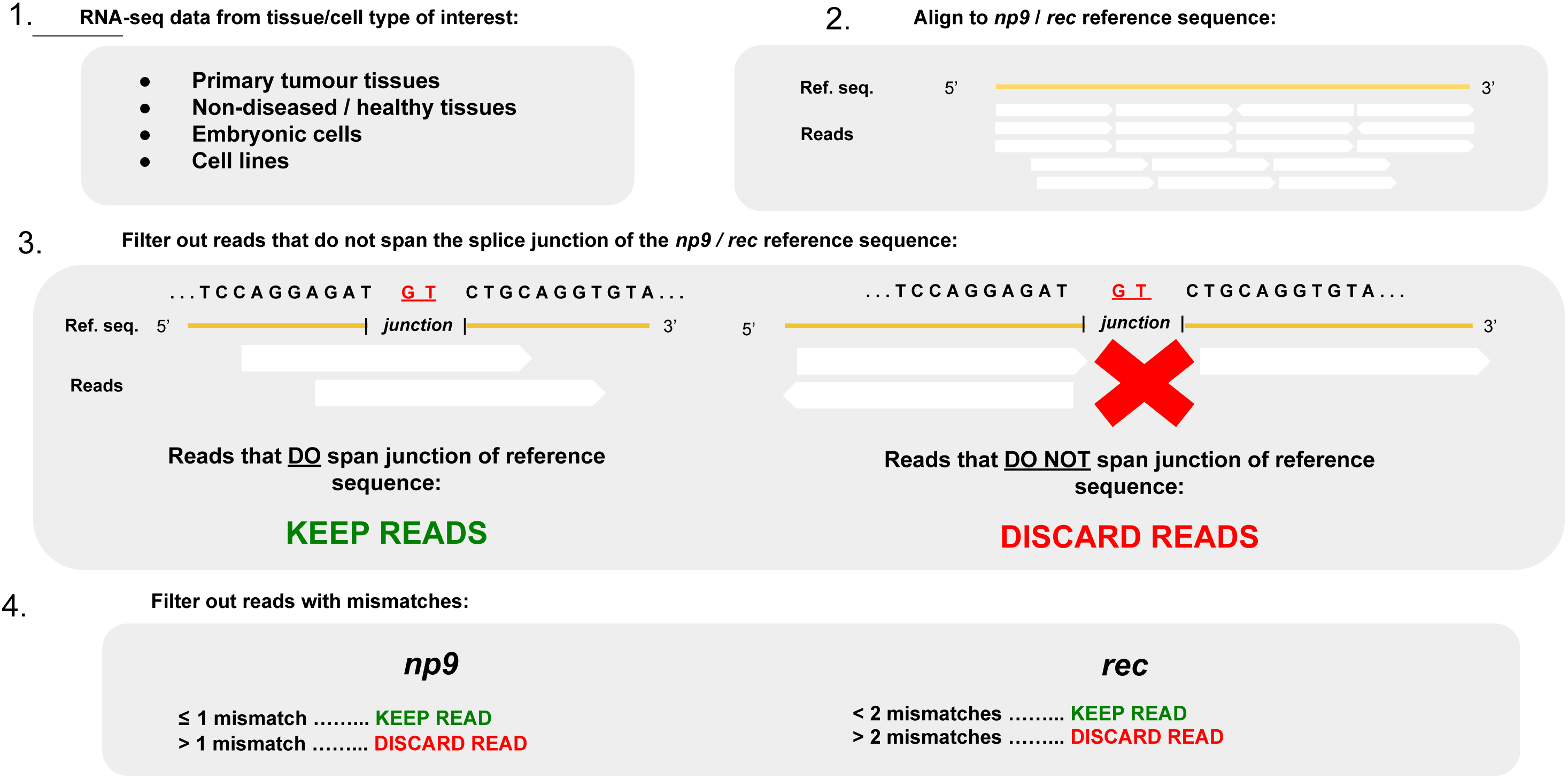
Workflow describing method in measuring expression of *np9* and *rec* in RNA-seq data. **1.** Download RNA-seq files from the tissue/cell type of interest. This can include transcriptional sequence data from primary tumour tissues, non-diseased or healthy tissues, embryonic cells, and cell lines. **2.** Align raw RNA-seq data to the *np9* or *rec* reference sequence (**Table S1**). **3.** Filter out reads that do not span the splice junctions of the *np9* /*rec* reference sequence, keeping only the reads that do span the respective splice junctions of *np9* and *rec*. **4.** Filter out the reads with sequence mismatches. For *np9*, if there’s more than 1 mismatch in these reads, discard the reads; for *rec,* if there’s more than 2 mismatches in these reads, discard the reads.

### Quantifying depth of coverage of HK2

The average depth of coverage of the mapped reads was quantified using the GenomeAnalysis TK package (GATK) version 3.6.0 [25] and Picard Tools version 2.1.1 (Broad Institute). For the other HK2 cDNA targets *pol* and LTR, as well as for the normalisation gene *actb*, the stringent filtering and processing necessary for capturing *np9* and *rec* transcripts was not necessary, and we used the average read count across the entire reference sequence. Because of this, the depth of coverage of *pol* and LTR may not be directly comparable to the depth of coverage observed for *np9* and *rec* expression, for each given sample. For these genes the BAM file with reads of a MAPQ >20 is directly input into Picard Tools for sorting and indexing.

### *Normalisation of HK2 transcripts by* actb *transcripts*

The read counts generated by GATK DepthOfCoverage were analysed in R version 3.3.1 [26]. Normalised expression of the HERV transcripts (*np9*, *rec*, LTR, and *pol*) from the samples was calculated by dividing the average depth of coverage of the expressed HERV by the average depth of coverage of the expressed reference gene (*actb*).

### Analysis of raw data

Raw FASTQ files of unaligned Illumina RNA-seq reads sequenced from 305 primary solid tumour tissues and blood-derived non-diseased tissues from different individuals were downloaded from Cancer Genomics Hub (data now available at Genomics Data Commons). Cancers that had been reported in the literature to be associated with HK2 expression were selected for study. The other cancer types were arbitrarily selected, and a total of 15 different cancers were analysed.

RNA-seq data sequenced from single cells of oocytes, zygotes, 2-, 4-, 8-cell embryos, morula, late blastocysts, and also early passages of human embryonic stem cell lines (71 files in total) were downloaded from Yan *et al.* [20] (**Table 1**). The full list of all of the files downloaded, including the accession and ID numbers are listed in Supplementary Materials.

**Table 1.** **Summary of RNA-seq data from solid primary tumour/non-diseased tissue**

| Tissue Source | Tissue Type |
| --- | --- |
| Breast invasive carcinoma (BRCA) | Primary solid tumour, solid tissue normal (NT) |
| Cervical squamous cell carcinoma/endocervical carcinoma (CESC) | Primary solid tumour, solid tissue normal (NT) |
| Cholangiocarcinoma (CHOL) | Primary solid tumour, solid tissue normal (NT) |
| Colon adenocarcinoma (COAD) | Primary solid tumour, solid tissue normal (NT) |
| Lymphoid neoplasm large B cell lymphoma (LBCL) | Primary solid tumour |
| Liver hepatocellular carcinoma (LIHC) | Primary solid tumour, solid tissue normal |
| Lung adenocarcinoma (LUAD) | Primary solid tumour, solid tissue normal (NT) |
| Lung squamous cell carcinoma (LUSC) | Primary solid tumour, solid tissue normal (NT) |
| Ovarian serous cystadenocarcinoma (OV) | Primary solid tumour |
| Pancreatic adenocarcinoma (PAAD) | Primary solid tumour, solid tissue normal |
| Prostate adenocarcinoma (PRAD) | Primary solid tumour, solid tissue normal (NT) |
| Skin cutaneous melanoma (SKCM) | Primary solid tumour, solid tissue normal (NT) |
| Testicular germ cell tumour (TGCT) | Primary solid tumour |
| Thyroid carcinoma (THCA) | Primary solid tumour, solid tissue normal (NT) |
| Uveal melanoma (UVM) | Primary solid tumour |
| Ovary* | Solid tissue normal (NT) |
| Testis* | Solid tissue normal (NT) |

All except for marked \* were sequenced at the University of North Carolina Lineberger Comprehensive Cancer Centre and come from The Cancer Genome Atlas project. Those marked with \* were sequenced at the European Bioinformatics Institute and come from the Illumina Human Body Map 2.0 Project.

### Non-diseased reference tissue selection

We first wanted to evaluate what the background expression of HK2 genes *np9* and *rec* are in normal, non-tumour tissues. Normal matched controls are not available for the majority of the tumour samples in TCGA so we decided to use a selection of random normal non-tumour tissues that are provided by TCGA and Illumina Human Body Map 2.0. The non-diseased tissue (NT) reference group is a mix of different healthy solid tissues, the majority of which are matched control tissue samples downloaded from TCGA. These tissue types include the cervix, bile duct, bowel, liver, lung, ovary, pancreas, prostate, testis, and thymus. To ensure that there is no significant bias in the non-diseased reference tissue samples selected for this comparison group, we also measured the expression of other groups of non-cancerous tissues from TCGA. These other groups (Table 2) consist of non-diseased breast tissue samples (BRCA_NT), tissues that came from organs that adenocarcinomas are derived from (AD_NT – bile duct, bowel, lung, pancreas, prostate), tissues that come from organs that squamous cell carcinomas are derived from (SQ_NT – cervix, lung), and tissues that come from organs that cancer types are derived from that do not fit in the other categories (OT_NT – liver, thymus). All accession/ID numbers of the samples used are in Supplementary Materials.

**Table 2.** **Summary of groups of non-diseased reference tissue samples used for analysis**

| Category | Tissue source |
| --- | --- |
| NT (non-diseased tissue reference group) | Bile duct ($n = 2$), cervix ($n = 3$), bowel ($n = 1$), liver ($n = 4$), lung ($n = 2$), ovary ($n = 1$), pancreas ($n = 1$), prostate ($n = 1$), testis ($n = 1$), thymus ($n = 2$) |
| BRCA_NT (breast tissue only) | Breast ($n = 18$) |
| AD_NT (organs that adenocarcinomas are derived from) | Bile duct (*n* = 5), bowel (*n* = 5), lung (*n* = 9), pancreas (*n* = 4), prostate (*n* = 5) |
| SQ_NT (organs that squamous cell carcinomas are derived from) | Cervix (*n* = 3), lung (*n* = 5) |
| OT_NT (organs that other cancer types are derived from) | Liver (*n* = 10), thymus (*n* = 5) |

### *Calculation of relative difference in expression between* np9 *and* rec *in cancers compared to non-diseased tissue reference group*

In order to determine the difference in relative expression of *np9* and *rec* in the different cancers compared to the non-diseased tissue reference group, the average read count of the HERV gene of each sample was divided by the average read count of *actb* within the same sample. The average normalised read count of HERV gene/*actb* of each sample is then calculated for each tissue group. Finally, the ratio of the average relative expression value of the HERV gene of each cancer type divided by the average relative expression value of the non-diseased tissue reference group (NT) is calculated.

### Statistical Analysis

The Wilcoxon signed rank-sum test was used to compare the relative expression (normalised within each sample using *actb*) of the HERV genes from each of the unmatched samples within each cancer type compared to the non-diseased tissue reference group (NT). The Wilcoxon signed rank-sum test was also used in comparing the relative expression of the same HK2 genes within the NT group to other non-diseased tissue categories (BRCA_NT, AD_NT, SQ_NT, OT_NT). Results with a p < 0.05 are considered significant. We also corrected for multiple comparisons using the Bonferroni method and report p-values < 0.0033 as significant.

For visualisation, the normalised values were transformed in order to account for samples with read counts of 0: log((normalised read count)*10,000+1),10). For all statistical methods, p-value < 0.05 is considered statistically significant. All statistical analyses were performed with R (version 3.3.1) in the integrated development environment (IDE) “Rstudio” (version 1.1.383)

## Results

### Describing HK2 expression in different categories of non-diseased tissue

To ensure that there is no significant bias in the non-diseased tissue samples selected for this comparison group, we also measured the expression of other groups of non-cancerous tissues from TCGA and found that there was no significant difference between the expression of the HERV targets in non-diseased tissue reference group (NT) and other non-diseased tissue types. *np9* and *rec* in the non-diseased tissue reference group (NT) is not significantly different compared to the expression of the same HERV genes in breast tissues (BRCA_NT), tissues from organs that adenocarcinomas are derived from (AD_NT), tissues from organs that squamous cell carcinomas are derived from (SQ_NT), and tissues from organs with cancer types that do not fit in the other categories (OT_NT) (**Fig. 4**, **Table 3**). *np9* and *rec* expression from BRCA_NT, AD_NT, SQ_NT, and OT_NT compared to NT was not statistically significantly different (p>0.05). However, a significant difference in relative expression of *pol* was observed in samples from the BRCA_NT group compared to NT (p = 0.007874). The expression of LTR was also significantly different within the NT samples compared to the AD_NT group (p = 0.01265) and SQ_NT (p = 0.045699). Because there appears to be no significant difference in expression levels of *rec* and *np9* between the different NT groups, we reasoned that the samples used in NT would be adequate to use as a non-diseased reference group when comparing expression profiles to different cancers.

**Figure 4.**
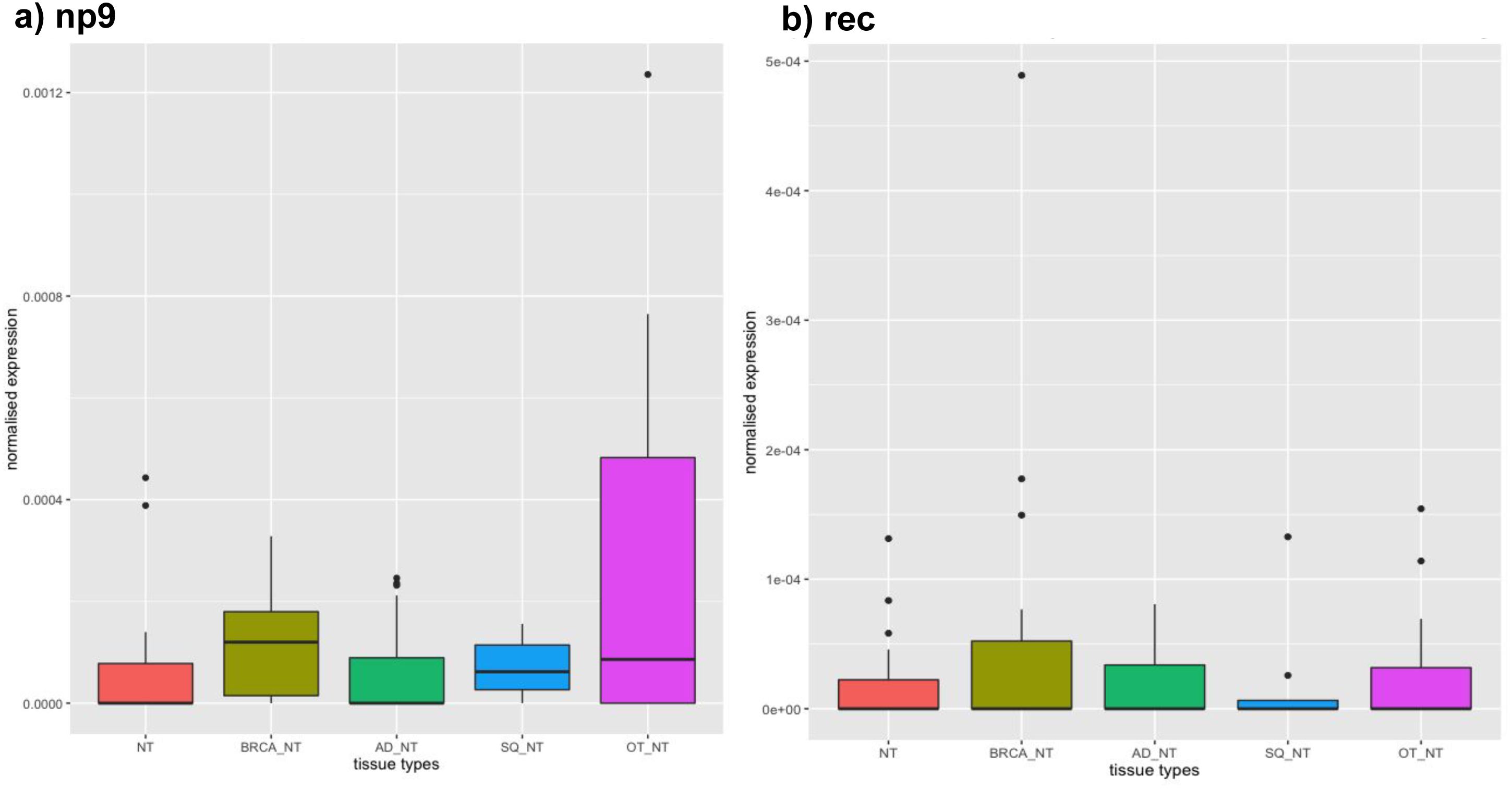
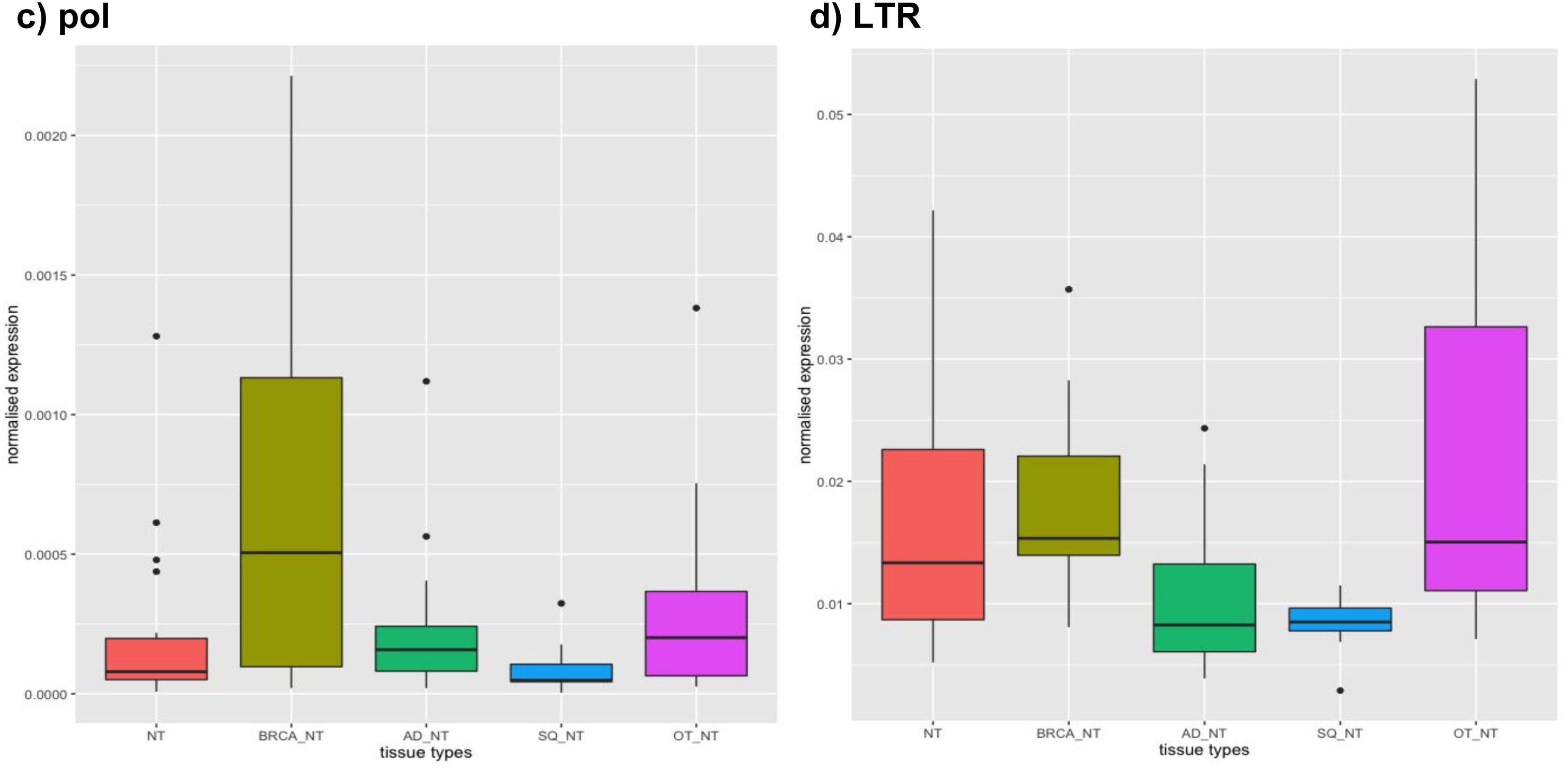
Describing HK2 expression in different categories of non-diseased tissue. x-axis indicates non-diseased tissue group types (from left to right): NT – pan-non-diseased tissue reference which includes tissues from bile duct, cervix, bowel, liver, lung, ovary, pancreas, prostate, testis, and thymus (*n* = 18), BRCA_NT – breast tissue (*n* = 18), AD_NT – tissues from bile duct, bowel, lung, pancreas, and prostate (*n* = 28), SQ _NT – tissues from cervix and lung (*n* = 8), OT_NT – tissues from liver and thymus (*n* = 15); y-axis indicates HERV transcription normalised by transcription of *actb*. Black horizontal lines indicate the mean. Non-diseased reference tissue (NT) group is placed on the far left in all box and whisker plots, for clarity. Shown is the relative expression of **(a.)** *np9*; **(b.)** *rec*; **(c.)** HK2 *pol*; **(d.)** HERVK-113 LTR across the different categories of non-diseased tissues.

**Table 3.** **Relative expression of HK2 in different non-diseased tissue group categories compared to non-diseased tissue reference (NT).**

**a)**
| *np9* |  |  | *rec* |  |  |
| --- | --- | --- | --- | --- | --- |
| Category | W test statistic | p-value | Category | W test statistic | p-value |
| BRCA_NT | 219 | 0.0635 | BRCA_NT | 181 | 0.496 |
| AD_NT | 247 | 0.9109 | AD_NT | 240 | 0.751 |
| SQ_NT | 90 | 0.306 | SQ_NT | 68 | 0.812 |
| OT_NT | 169 | 0.1925 | OT_NT | 133 | 0.937 |

**b)**
| *pol* |  |  | LTR |  |  |
| --- | --- | --- | --- | --- | --- |
| Category | W test statistic | p-value | Category | W test statistic | p-value |
| BRCA_NT | 245 | 0.007874 * | BRCA_NT | 190 | 0.389 |
| AD_NT | 307 | 0.2224 | AD_NT | 142 | 0.0127 * |
| SQ_NT | 54 | 0.3383 | SQ_NT | 36 | 0.0457 * |
| OT_NT | 162 | 0.3428 | OT_NT | 160 | 0.381 |

**(a.)** The expression of *np9* and *rec* and the expression of **(b.)** HK2 *pol* and HERVK-113 LTR in samples within the NT reference group in comparison to other non-diseased tissue groups.
\* indicate *p* < 0.05

### HK2 expression in early embryogenesis and stem cell lines

Transcription of *np9* and *rec* is activated at, and is highest at, the 8-cell stage of early embryogenesis, but *rec* expression is overall silenced during the following developmental stages (**Fig. 3b**). Interestingly, *np9* is transcriptionally active (but downregulated) in morula and in the early passages of the hESC lines (**Fig. 3a**), and both *pol* and HERV-K113 LTR show upregulation in the late blastocyst that is not seen in *np9* and *rec* (**Fig. 3c-d**).

**Figure 3.**
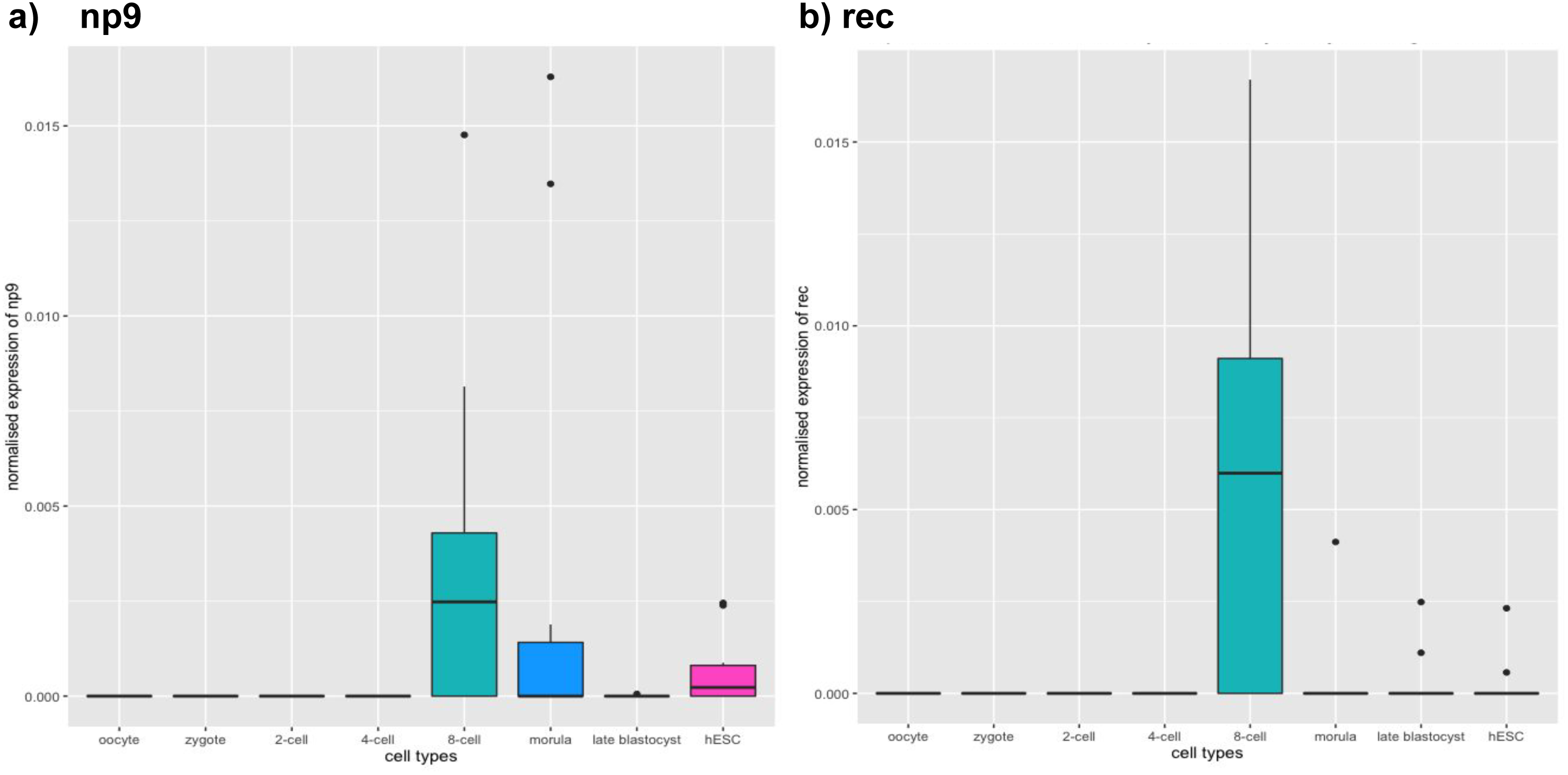
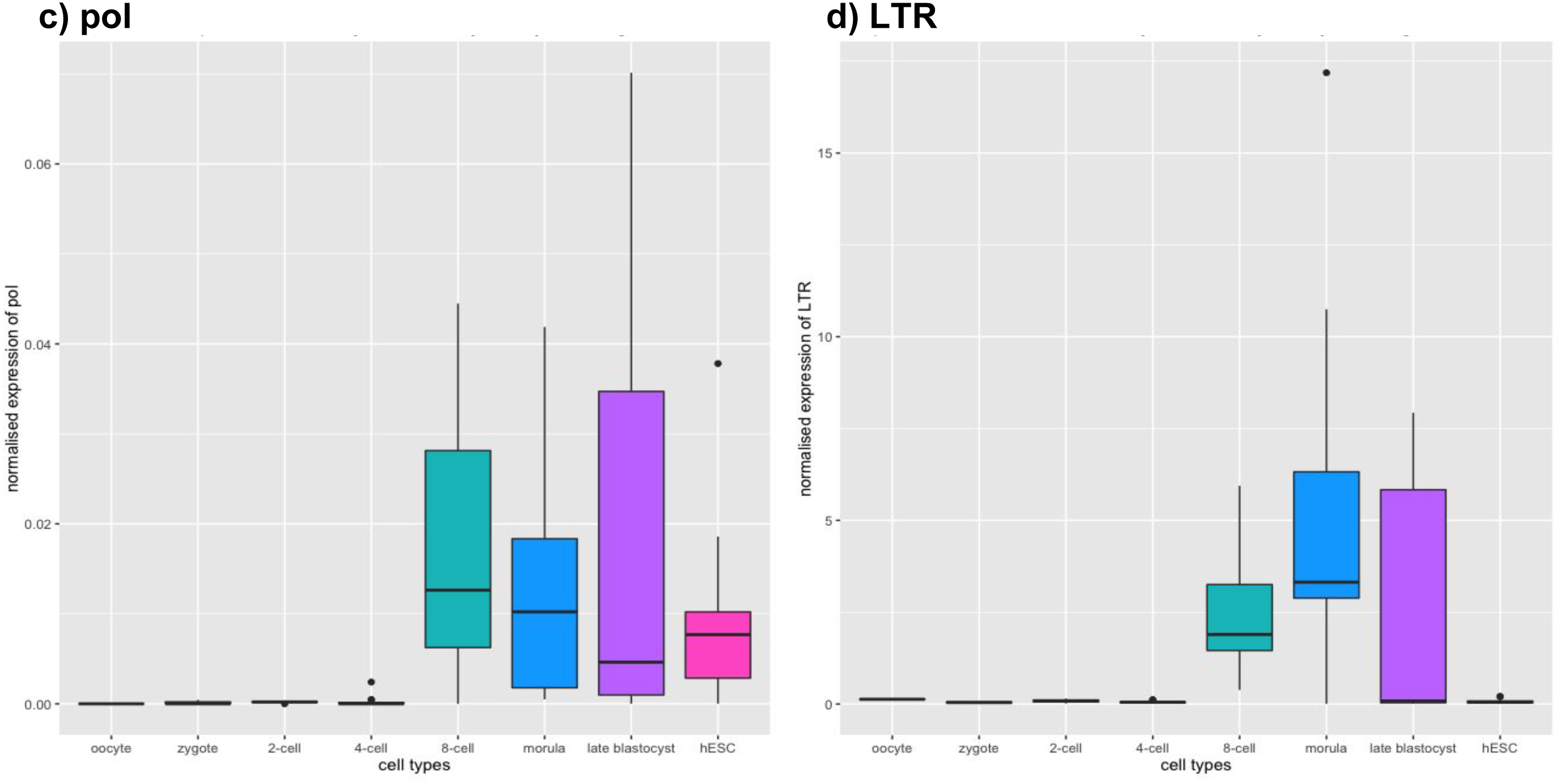
**Expression of HK2 targets in human preimplantation embryos and embryonic stem cells.** x-axis indicates cell types (from left to right): Oocyte (*n =* 3), zygote (*n* = 3), 2-cell (*n* = 4), 4-cell (*n* = 12), 8-cell (*n* = 17), morula (*n* = 10), late blastocyst (*n* = 10), hESC (*n* = 13); y-axis indicates HERV transcription normalised by transcription of *actb*. Black horizontal lines indicate the median. Data are from (15). Expression of **(a.)** *np9*; **(b.)** *rec*; **(c.)** HK2 *pol*; **(d.)** HERVK-113 LTR in the different stages of human preimplantation embryos and human embryonic stem cell lines.

### HK2 expression in non-diseased tissues and cancer samples

When we compared the expression of the HK2 genes *np9* and *rec* in non-diseased tissue reference and the 15 different cancers, we found that there is a general trend of HERV gene upregulation across the cancers, with some cancers that show significant overexpression **(Table 4)**. *np9* is overexpressed by at least 2x higher in 9 out of 15 cancers compared to the non-diseased tissue reference group, with significant upregulation by almost 5x in cholangiocarcinoma (p = 0.009), almost 5.5x higher expression in liver hepatocellular carcinoma (p = 0.006), 4x higher expression in lung adenocarcinoma (p = 0.001), almost 2x higher expression in thyroid carcinoma (p = 0.043), and nearly 13x higher expression in uveal melanoma (p = 0.011). *np9* was highly upregulated in testicular germ cell cancer, with 55.5x fold change in overexpression compared to the non-diseased tissue group testicular germ cell cancer (p = 9.83 x 10^-7^) **(Fig. 5a)**. On the other hand, *rec* is overexpressed compared to the non-diseased tissue reference group in 6 out of the 15 cancers surveyed, and significantly upregulated in 3 cancers. In skin cutaneous melanoma, *rec* is overexpressed by almost 44 times compared to the expression of *rec* in the non-diseased tissue group (p = 0.041), overexpressed by 83x in testicular germ cell cancer (p = 1.53 x 10^-6^), and is highly upregulated in uveal melanoma by more than a thousandfold (p = 5.81 x 10^-7^) (**Fig. 5b**).

**Figure 5.**
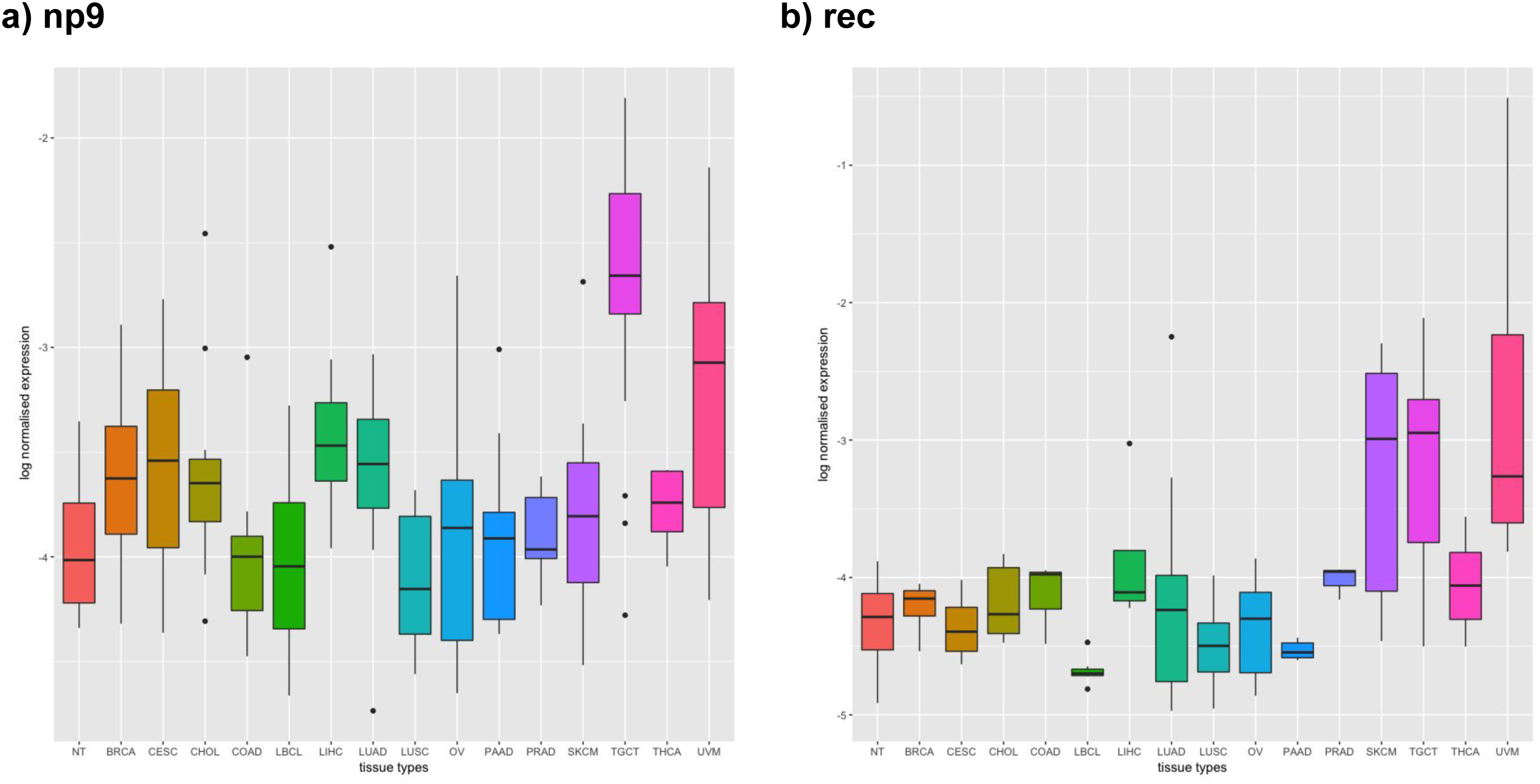
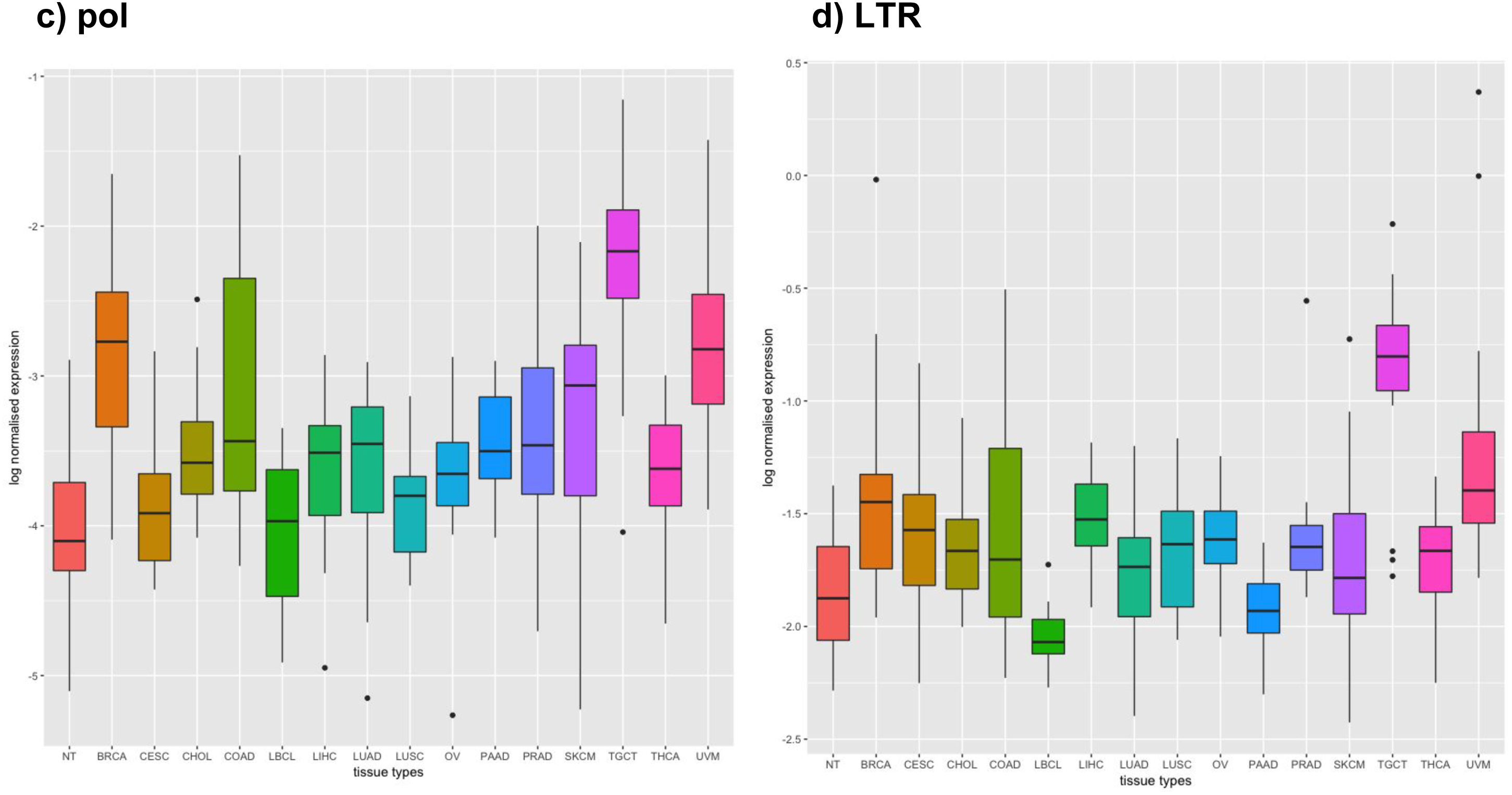
**Expression of HK2 genes in non-diseased reference tissue group (NT) compared to 15 different cancers**. x-axis indicates tissue types (from left to right): NT – normal tissue reference group (*n* = 18), BRCA – breast invasive carcinoma (*n* = 20), CESC – cervical squamous cell carcinoma/endocervical adenocarcinoma (*n* = 18), CHOL – cholangiocarcinoma (*n* = 20), COAD – colorectal adenocarcinoma (*n* = 20), LUSC – lung squamous cell carcinoma (*n* = 20), OV – ovarian serous cystadenocarcinoma (*n* = 20), PAAD – pancreatic adenocarcinoma (*n* = 20), PRAD – prostate adenocarcinoma (*n* = 20), SKCM – skin cutaneous melanoma (*n* = 24), TGCT – testicular germ cell cancer (*n* = 20), THCA – thyroid carcinoma (*n* = 20), UVM – uveal melanoma (*n* = 20); y-axis indicates HK2 transcription normalised by transcription of *actb*, on a scale transformation of log_10_(normalised read count)*10,000+1). Black horizontal lines indicate the median. Normal tissue (NT) group is placed on the far left for clarity. Expression of **(a.)** *np9*; **(b.)** *rec*; HK2 *pol*; and **(d.)** HERVK-113 LTR in NT compared to 15 different cancers.

**Table 4.**
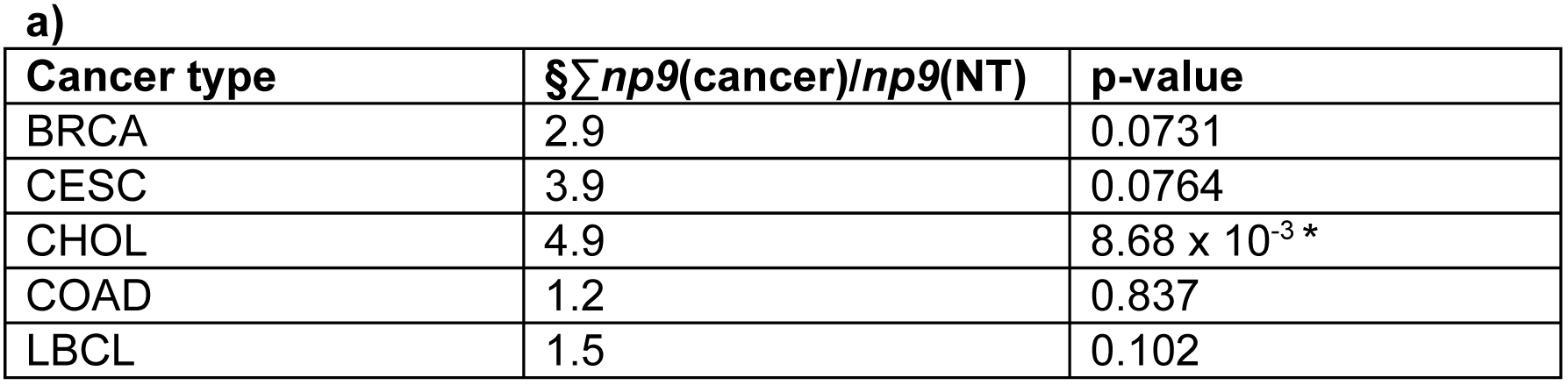
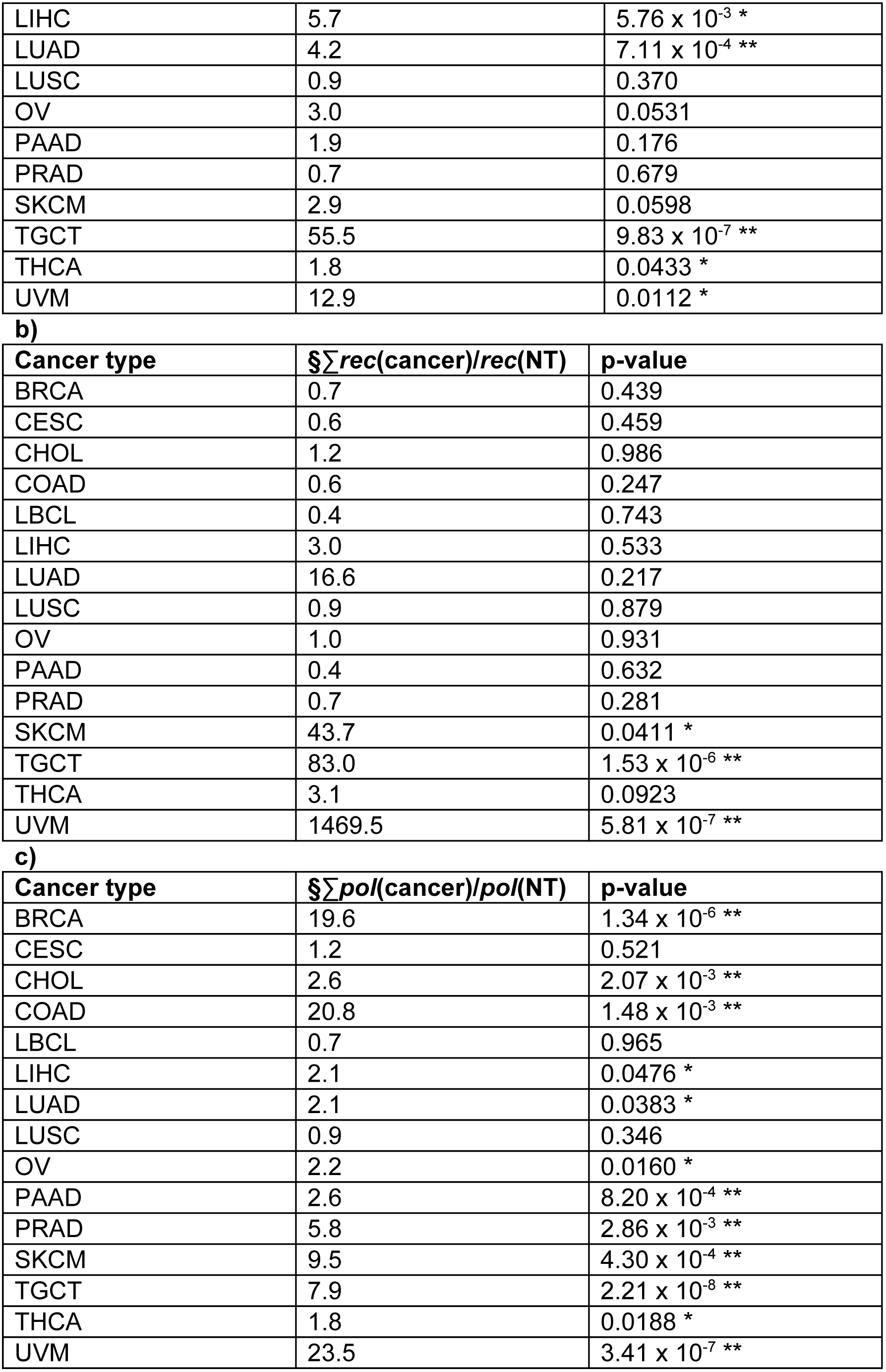
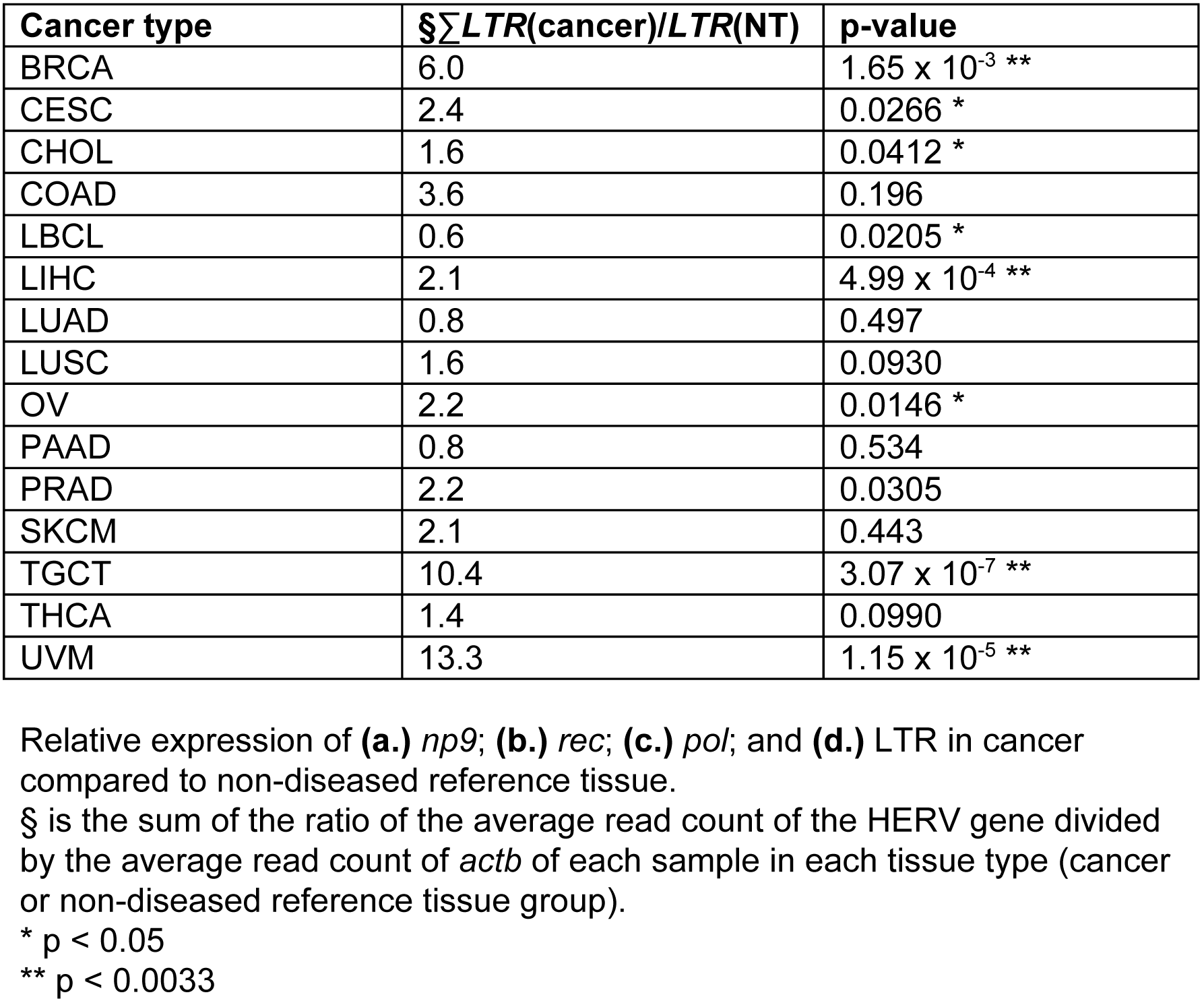
**Relative expression of HK2 targets in cancers compared to NT**

**a)**
| Cancer type | $\sum np9(\text{cancer})/np9(\text{NT})$ | p-value |
| --- | --- | --- |
| BRCA | 2.9 | 0.0731 |
| CESC | 3.9 | 0.0764 |
| CHOL | 4.9 | $8.68 \times 10^{-3} *$ |
| COAD | 1.2 | 0.837 |
| LBCL | 1.5 | 0.102 |
| LIHC | 5.7 | $5.76 \times 10^{-3} *$ |
| LUAD | 4.2 | $7.11 \times 10^{-4} **$ |
| LUSC | 0.9 | 0.370 |
| OV | 3.0 | 0.0531 |
| PAAD | 1.9 | 0.176 |
| PRAD | 0.7 | 0.679 |
| SKCM | 2.9 | 0.0598 |
| TGCT | 55.5 | $9.83 \times 10^{-7} **$ |
| THCA | 1.8 | $0.0433 *$ |
| UVM | 12.9 | $0.0112 *$ |

b)
| Cancer type | $\sum \text{rec}(\text{cancer})/\text{rec}(\text{NT})$ | p-value |
| --- | --- | --- |
| BRCA | 0.7 | 0.439 |
| CESC | 0.6 | 0.459 |
| CHOL | 1.2 | 0.986 |
| COAD | 0.6 | 0.247 |
| LBCL | 0.4 | 0.743 |
| LIHC | 3.0 | 0.533 |
| LUAD | 16.6 | 0.217 |
| LUSC | 0.9 | 0.879 |
| OV | 1.0 | 0.931 |
| PAAD | 0.4 | 0.632 |
| PRAD | 0.7 | 0.281 |
| SKCM | 43.7 | $0.0411 *$ |
| TGCT | 83.0 | $1.53 \times 10^{-6} **$ |
| THCA | 3.1 | 0.0923 |
| UVM | 1469.5 | $5.81 \times 10^{-7} **$ |

c)
| Cancer type | $\sum \text{pol}(\text{cancer})/\text{pol}(\text{NT})$ | p-value |
| --- | --- | --- |
| BRCA | 19.6 | $1.34 \times 10^{-6} **$ |
| CESC | 1.2 | 0.521 |
| CHOL | 2.6 | $2.07 \times 10^{-3} **$ |
| COAD | 20.8 | $1.48 \times 10^{-3} **$ |
| LBCL | 0.7 | 0.965 |
| LIHC | 2.1 | $0.0476 *$ |
| LUAD | 2.1 | $0.0383 *$ |
| LUSC | 0.9 | 0.346 |
| OV | 2.2 | $0.0160 *$ |
| PAAD | 2.6 | $8.20 \times 10^{-4} **$ |
| PRAD | 5.8 | $2.86 \times 10^{-3} **$ |
| SKCM | 9.5 | $4.30 \times 10^{-4} **$ |
| TGCT | 7.9 | $2.21 \times 10^{-8} **$ |
| THCA | 1.8 | $0.0188 *$ |
| UVM | 23.5 | $3.41 \times 10^{-7} **$ |

d)
| Cancer type | $\sum LTR(\text{cancer})/LTR(\text{NT})$ | p-value |
| --- | --- | --- |
| BRCA | 6.0 | $1.65 \times 10^{-3}$ ** |
| CESC | 2.4 | 0.0266 * |
| CHOL | 1.6 | 0.0412 * |
| COAD | 3.6 | 0.196 |
| LBCL | 0.6 | 0.0205 * |
| LIHC | 2.1 | $4.99 \times 10^{-4}$ ** |
| LUAD | 0.8 | 0.497 |
| LUSC | 1.6 | 0.0930 |
| OV | 2.2 | 0.0146 * |
| PAAD | 0.8 | 0.534 |
| PRAD | 2.2 | 0.0305 |
| SKCM | 2.1 | 0.443 |
| TGCT | 10.4 | $3.07 \times 10^{-7}$ ** |
| THCA | 1.4 | 0.0990 |
| UVM | 13.3 | $1.15 \times 10^{-5}$ ** |

Relative expression of **(a.)** *np9*; **(b.)** *rec*; **(c.)** *pol*; and **(d.)** LTR in cancer compared to non-diseased reference tissue.
§ is the sum of the ratio of the average read count of the HERV gene divided by the average read count of *actb* of each sample in each tissue type (cancer or non-diseased reference tissue group).
\* $p < 0.05$
\*\* $p < 0.0033$

## Discussion

Retroviruses have the ability to contribute to tumourigenesis by activating cellular oncogenes or mutating cellular tumour suppressor genes, resulting in aberrant gene expression that promotes a cascade of uncontrolled cellular proliferation [27],[28]. Although these tumourigenic mechanisms by ERVs have been observed in other host organisms, namely in mice, there is yet no evidence of ERVs directly contributing to cancer pathogenesis in humans. In comparison to non-diseased tissue, we had expected to see HK2 target overexpression in breast cancer, germ cell tumours, skin melanoma, lymphoma, prostate cancer, liver cancer, and prostate cancer. We also surveyed other cancer types that, to our knowledge, have not been examined for HK2 expression. In addition, because the HK2-encoded accessory genes *np9* and *rec* may be considered as biomarkers to specific cancers, due to their association with the activation of signalling pathways implicated in cancers, it was our objective to explore their tissue-specific expression further.

In agreement with previously reported results [7],[29], HK2 appears to be derepressed and transcriptionally active starting from the 8-cell and 16-cell stage (morula) of human embryonic development, and varying levels of transcription in the blastocyst and in its outgrowths.

Breast cancer and testicular germ cell tumours are all cancers that derive from organs of the endocrine system. It is tempting to speculate that this link between HERV-K expression and some these cancer types is due to the expression of steroid hormones in glandular tissue. For example, within the U3 region of the HERV-K LTR, there are several predicted oestrogen, progesterone, and androgen response elements. *env* transcription has been shown to be upregulated up to 10-fold in breast cancer cell lines that had been treated with oestradiol followed by progesterone. Moreover, HERV-K reverse transcriptase is overexpressed in breast cancer cell lines that have been treated with the oestradiol/progesterone combination, suggesting that these steroid hormones indeed have a regulatory effect on HERV transcription [30],[31].

Because of the novel method we developed in processing and analysing the expression of highly similar repetitive elements in RNA-seq data, the strict parameters we used in interpreting the significance of the results may have led to false negatives. The Wilcoxon signed rank-sum test was chosen to compare the relative expression of the HK2 genes from each of the unmatched samples within each cancer type compared to non-diseased tissue, and the values were corrected for multiple comparisons using the Bonferroni method. p-values < 0.0033 were considered significant. Although, according to our parameters of significance, only a few cancers were determined as having *np9*, and *rec* transcripts upregulated in comparison to non-diseased tissue (**Fig. 2**, **Table 4**), there is a general trend of HERV upregulation in the majority of the cancers surveyed, in comparison to the median expression of non-diseased tissue, including the cancer types that did not met our threshold of significance. Thus, the lack of significance in correlations and comparisons is likely to reflect lack of power by adopting a conservative p-value rather than true absence of correlation.

HERV expression is very well-documented in testicular germ cell tumours – germ cell tumour cell lines have been shown to express HK2 non-infectious particles [32],[33],[34]. HERV-K has also been shown to be overexpressed at specific stages of early embryonic development[7],[9],[11], which we also observed using our novel method. Germ cell tumours also share with early embryonic cells similar traits of embryonic stem cell identity, such as the upregulation of the pluripotency factors Oct3/4, Nanog, and Sox2 [35].

During the early stages of normal development, primordial germ cells are also totipotent stem cells: the expression of pluripotency factors Oct3/4, Nanog, Sox2 gives primordial germ cells high self-renewal capacity by the suppression of differentiation [35]. A characteristic chromosomal abnormality seen in testicular germ cell tumours is a gain of the short arm of chromosome 12 (chr12p), resulting in the amplification of genes that map to that location, which includes stem-cell associated factors like Nanog, cell cycle regulators like cyclin D2 (*CCND2*), the oncogene KRAS, the growth factor receptor tumour necrosis factor receptor 1 (*TNFRSF1A*), and genes involved in metabolism such as GAPDH and GLUT3 [35]. Oct4 and Nanog are known to bind to HERV LTRs, mediating the transcription of both host genes and the provirus [7]. It has been suggested that the increased levels of Nanog due to the chr12p gain may be enough to overexpress the other core pluripotency factors Oct3 and Sox2, encoded by genes located on different chromosomes[35].

We observed a bias in the abundance of transcription from the 5’ end of HERV-K113 LTR, where the U3 region is (**Fig. 2a**), compared to the 3’ end. In particular, transcription is most abundant at the GA-rich motif found at the 5’ end. The uneven distribution of mapped reads could be due to biological biases in provirus expression. For example, in teratocarcinoma cell lines, both LTRs of full proviruses are transcribed, although the 3’ LTR appears to be transcribed far less abundantly than the 5’ LTR [36]. However, the significance or functional importance of truncated expression from the 3’ LTR is unknown. The formation of cancers often does not have a single aetiological agent that directly forms the disease. Tumourigenesis and the formation of malignancies is a complex, multifactorial process that requires the presence of both genetic predisposition of the individual to the cancer as well as life history factors such as environmental exposure to carcinogens. It is possible that the polymorphic nature of HK2 as well as its potential to express oncogenes like *np9* and *rec* are key components in increasing the genetic susceptibility of an individual to that cancer.

It would also be interesting to further explore the scope of *np9* and *rec* upregulation in even more cancers. Although we used RNA-seq files from 15 different types of cancer, there are more cancer types reported in the literature to overexpress HERVs: colorectal cancer [37], chronic lymphocytic leukaemia (CLL) [38], and Burkitt’s lymphoma [39]. Further research could also be performed to characterise the broad network that indicates how Np9 and Rec proteins interact with effectors in the MAPK/Akt/NOTCH signalling cascade that ultimately leads to cell survival and proliferation, specific to the cancers they are upregulated in.

Limitations: The method for quantifying HK2 transcripts [17] is under validation and the results should be considered as preliminary/pending verification from a validation process.

## Acknowledgements

Thank you to Fabricia Nascimento, Robert Belshaw, Tim Coulson, and the Palaeovirology group for helpful discussions and comments.

## Ethics Approval and Consent to Participate

All human data used were obtained through The Cancer Genome Atlas and used according to the Data Use Certification as stipulated by the TCGA Data Access Committee and dbGaP authorisation (https://dbgap.ncbi.nlm.nih.gov/aa). Permission to use controlled-access data was approved by dbGaP under projects #57108 and #7621.

